# Vibe Coding Specificity Foundation Models

**DOI:** 10.64898/2026.06.04.730134

**Authors:** Sai T. Reddy

## Abstract

Molecular recognition — the determination of which agent binds which target — governs adaptive immunity, gene regulation, signal transduction, RNA silencing, enzyme catalysis, and the selectivity of therapeutics. Determining binding specificity remains dependent on experimental screening or domain-specific computational tools that do not generalize across binding modalities. Transformer softmax attention is mathematically identical to the Boltzmann distribution governing molecular binding^1^. This identity, together with five conditions of molecular recognition systems, prescribes a single neural network architecture for cross-modal binding prediction: dual sequence encoders, symmetric contrastive learning, and a learned physical temperature^2^. A Specificity Foundation Model (SFM) is an instance of this physics-derived, sequence-to-sequence architecture that maps any agent–target sequence pair to a binding compatibility score, enabling bidirectional retrieval across molecular recognition domains without requiring structural information. The first SFM for antibody–antigen binding demonstrated ∼100,000-fold greater data efficiency than comparable vision-language models^3^. Here we report six SFMs across six molecular recognition domains — transcription factor–DNA, enzyme–substrate, peptide–MHC, CRISPR gRNA–off-target genomic DNA, microRNA–mRNA target, and small molecule drug–target protein — using the identical architecture without modification and trained using publicly available data only. Evaluated by cross-modal retrieval from pools of 512 candidates (random baseline 0.2%), in-distribution R@1 ranges from 27.7% to 98.0% across the six domains. mir-SFM retrieves miRNA targets at 98.0% R@1, including the ∼80% of validated interactions that seed-matching tools cannot find. mhcSFM achieves 95.4% R@1 on held-out rare HLA alleles absent from training. Applying crisprSFM to CRISPR off-target prediction improves precision to 94.0% compared to 33.2% from Hamming distance alone. All six SFMs were built by a domain expert with no programming experience using vibe coding — natural-language-directed AI coding agents — with numerical claims independently verified by an orthogonal AI auditor. These results establish SFMs as a physics-derived, sequence-native class of model that augments experimental and computational workflows across molecular recognition domains.

## INTRODUCTION

Molecular recognition determines which molecular agent binds which target, and with what specificity and is the organizing computation of biology. It governs adaptive immunity, gene regulation, signal transduction, RNA silencing, and the selectivity of small-molecule therapeutics. Predicting molecular recognition from sequence is the shared computational problem underlying antibody engineering, vaccine design, drug safety profiling, gene therapy target selection, and functional genomics. Despite this centrality, the reliable determination of binding specificity in nearly every domain still requires experimental screening: ChIP-seq and high-throughput SELEX for transcription factor–DNA recognition^4,5^, mass spectrometry immunopeptidomics for HLA peptide presentation^6^, GUIDE-seq, CIRCLE-seq and Discover-seq for CRISPR off-target cleavage^7–9^, AGO-CLIP-seq for miRNA target identification^10^, chemoproteomic and kinase-panel screens for small-molecule target selectivity^11^, and biochemical characterization for enzyme–substrate annotation^12^. These campaigns are costly, time-intensive, scale linearly with experimental capacity, and form the rate-limiting step of every downstream application that depends on binding specificity.

Computational tools exist for each of these domains but are often narrow sequence-based classifiers — models trained on domain-specific features, for specific targets or agent classes, that do not generalize beyond the task they were built for. NetMHCpan 4.1 predicts peptide–MHC binding for a fixed set of alleles^13^; TargetScan predicts canonical miRNA seed-matches^14^; Cas-OFFinder enumerates CRISPR off-targets by mismatch count^15^; DeepBind and BPNet predict transcription factor binding for individual TFs^16,17^. Each is a separate engineering effort with a domain-specific architecture; none is a foundation model in the sense of a single pretrained system that transfers across tasks and domains. Structure-based AI methods — AlphaFold 3^18^, Boltz-2^19^, RFdiffusion^20,21^, BindCraft^22^ — predict 3D complex geometry and have transformed protein structure prediction and de novo binder design^23,24^. These models incorporate physics as inductive bias but are not derived from a governing equation of molecular binding. Single-domain contrastive dual-encoder models — DrugCLIP^25^, ConPLex^26^, CLIPZyme^27^ — share architectural elements with the models reported here but were developed as bespoke solutions for individual domains and have not been demonstrated to transfer. None of the above answers the cross-modal retrieval question: given an agent, retrieve its target from a candidate pool by learned binding compatibility, and vice versa.

The softmax function in transformer attention is mathematically identical to the Boltzmann distribution governing molecular binding at thermal equilibrium^1^. The identity follows from the Luce uniqueness theorem (Luce, 1959)^28^: softmax attention is the unique probability rule consistent with Boltzmann-distributed scores under stochastic competitive binding. From this identity and five conditions of the molecular system — discrete intermolecular contacts, bilinear energy decomposition, finite competitor pools, thermal equilibrium, and stochastic selection — the architecture for any binding domain is prescribed: dual encoders mapping agent and target sequences into a shared embedding space, symmetric InfoNCE training, mean-pooled embeddings, and a learned temperature corresponding to physical *k*_*B*_*T*. We define a Specificity Foundation Model (SFM) as an instance of this physics-derived architecture^2^. Our recent work describes the Cross-attention Adaptive immune receptor-Antigen Language Model (CALM)^3^ as the first implementation of an SFM for antibody–antigen specificity, achieving cross-modal retrieval from ∼4,000 training pairs with approximately 100,000-fold greater data efficiency than contrastive learning models using text-image pairs^29^. Both predictions of the Molecular Recognition Computing (MRC) framework described in our previous work^2^ — that the architecture transfers across binding domains, and that training exhibits data efficiency consistent with parameter estimation in a known equation rather than function approximation — have been tested in this single domain of antibody-antigen specificity.

The SFM architecture differs from structure-based methods in a respect that follows directly from the governing equation. Structure-based models predict the three-dimensional geometry of a molecular complex given its components — the output is a structure, and the input typically requires or benefits from structural information. These models incorporate physics through inductive biases — SE(3) equivariance, atomic interaction potentials, coevolutionary signal — that constrain the model toward physically plausible outputs, a design philosophy the MRC framework terms Level 2 alignment^2^.

SFMs operate at Level 3: the architecture itself is derived from the governing equation of binding: the Boltzmann distribution (Second Law of Thermodynamics), which specifies that the probability of binding between agent *i* and target *j* is *P*(*j*|*i*) = exp(*s*_*ij*_*/τ*)*/* ∑_*k*_ exp (*s*_*ik*_*/τ*), where *s*_*ij*_ is the compatibility score and *τ* is temperature. This is a sequence-to-score computation: it requires only the sequences of the two binding partners, maps them to compatibility scores via learned embeddings, and produces a probability distribution over all candidate targets. No structural information is required for inference. In practical terms, a trained SFM accepts any agent sequence and any target sequence, computes binding compatibility in a single forward pass, and returns a ranked list of candidates in either direction. Inference is computationally light: a pool of 10,000 candidates can be scored in seconds on a single GPU, enabling high-throughput virtual screening at a scale and speed that experimental methods cannot match and that per-task classifiers were not designed to provide.

Vibe coding — iterative, natural-language-directed software construction using AI coding agents — is a methodology that has been applied across software engineering domains^30^ and is emerging as a tool for biological data analysis and scientific software^31^. A capability of AI coding agents that is particularly relevant for computational biology is the management of heterogeneous data formats, database APIs, file-format conversions, and preprocessing pipelines — tasks that constitute the most labor-intensive component of building any bioinformatics tool and that coding agents handle as routine operations. This paper reports its application to biological foundation-model construction. The scope is research prototypes, not production software. Vibe coding is tractable for SFMs because the physics-derived architectural prescription removes the architecture-search, loss-design, and ablation work that ML engineering typically supplies; what remains is scientific judgment — encoder selection, data curation, hard-negative strategy, evaluation design — and domain-expertise decisions such as the HLA binding-groove pseudo-sequence extraction used in mhcSFM. An orthogonal AI verification protocol, in which a second independent AI agent audits every numerical claim from committed repository artifacts, provides the verification layer.

Here, we report six SFMs across six molecular recognition domains, each representing the first SFM for its domain: transcription factor–DNA (tSFM), enzyme–substrate (eSFM), peptide–MHC class I (mhcSFM), CRISPR guide RNA–genomic off-target DNA (crisprSFM), microRNA–mRNA target (mir-SFM), and small molecule drug–target protein binding (dtSFM). The architecture, training objective, and evaluation protocol were unchanged across all six. All SFMs were built by a single domain expert with no prior coding experience, working with a directed AI coding assistant and the published CALM reference implementation, using only training data available from publicly available repositories and in a development window of weeks per SFM. The performance from retrieval pools of 512 candidates (random baseline 0.2%) of several prototypes is striking. mir-SFM retrieves miRNA targets at 98.0% R@1 — including the ∼80% of validated interactions that seed-based tools cannot find^32^. mhcSFM achieves 95.4% R@1 on held-out rare HLA alleles absent from training, and a cascade filter with NetMHCpan 4.1 nearly doubles MS-presentation precision from a cancer neoantigen benchmark^33^. crisprSFM paired with Hamming distance pre-filtering raises off-target precision from 33.2% to 94.0% at the clinical ≤4-mismatch threshold. dtSFM achieves 91.2% R@10 at 80% protein-identity out-of-distribution (OOD), with retrieval exceeding in-distribution, a pattern predicted by the MRC framework^2^ but absent from standard supervised learning. These prototype SFMs augment the experimental and computational tools each field already uses; as open-source, computationally lightweight, sequence-to-sequence systems, they can be deployed alongside existing workflows at minimal cost. Across the six domains, training dynamics — fast convergence, inverted train-test accuracy, OOD non-degradation — appear in four of six domains, consistent with the MRC framework’s prediction that SFM training is parameter estimation in a known governing equation rather than function approximation of an unknown function class. Each prototype admits a scaling path through additional paired-interaction data that exists today in public databases, encoder upgrades, and the cross-attentive decoder extension^2^ that converts every SFM into a bidirectional generative model capable of designing binding agents for specified targets. The six SFMs reported here are the empirical companion to our recent studies^1,2^: the demonstration that the convergence between softmax attention and Boltzmann-distributed binding produces working models across molecular recognition domains.

## RESULTS

### Vibe coding SFMs: physics-derived architecture enables domain-expert construction

The SFM architecture is prescribed by the convergence of softmax attention and Boltzmann distribution, what we refer to as the convergence equation, as well as five conditions of molecular recognition systems^1,2^ (**Fig. 1**). Because the architecture, loss function, and training protocol are fully specified by the physics, what remains for the domain expert is not ML engineering but scientific judgment: which paired-interaction database to use, which pretrained encoders match the molecular modality, what constitutes a biologically meaningful hard negative, and how to validate the result. This is the property that makes SFMs accessible to vibe coding.

**Figure 1.**
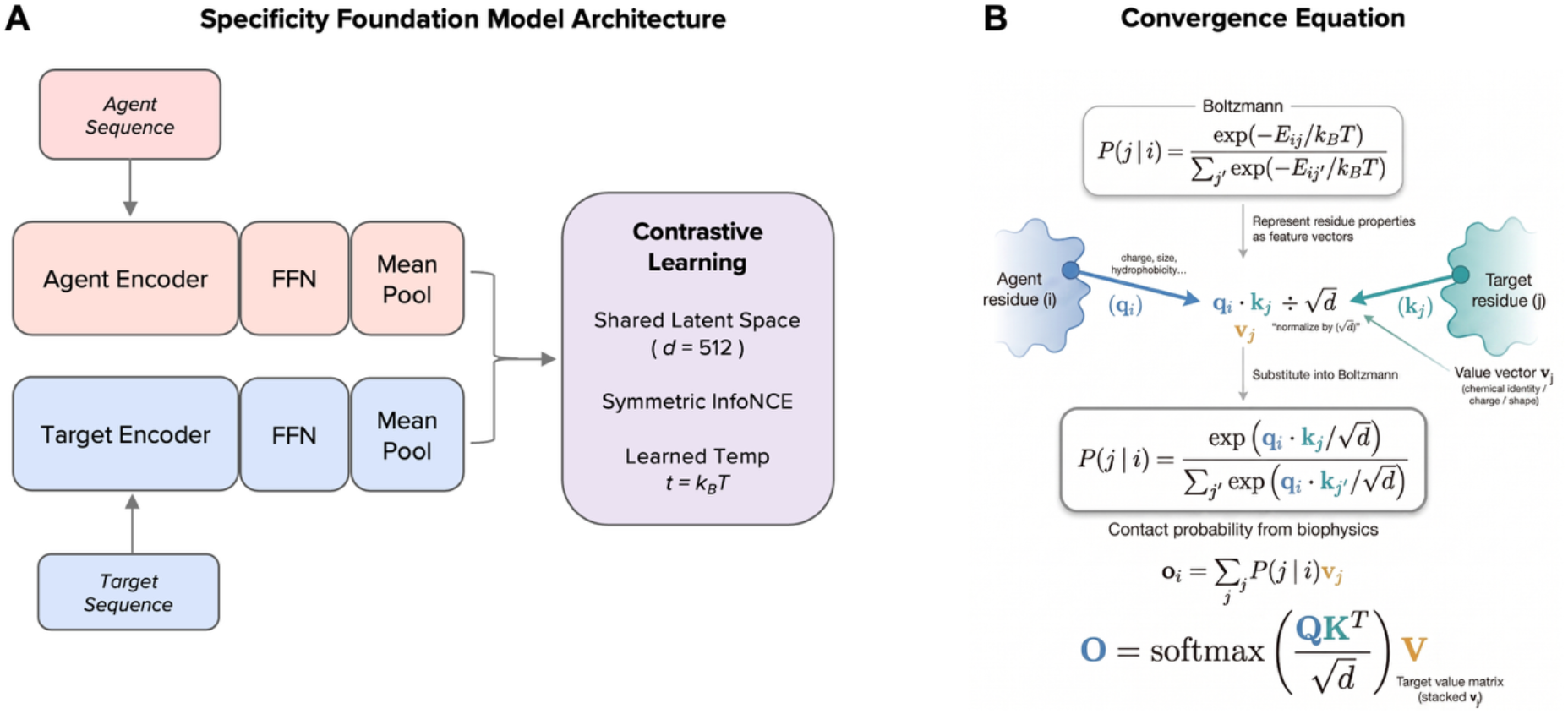
SFM architecture and the convergence equation. (A) The Specificity Foundation Model (SFM) architecture prescribed by the convergence equation and five conditions of molecular recognition^1,2^. Two pretrained sequence encoders (agent and target) project through feedforward networks (FFN) with mean pooling into a shared 512-dimensional latent space. Training minimizes the symmetric InfoNCE loss with a learned temperature parameter *τ* corresponding to physical *k*_*B*_*T*. Encoder weights are frozen; only the FFN projection heads (∼5–7 million parameters) are trained. The architecture is applied unchanged across all six molecular recognition domains reported in this paper; only the encoders, paired-interaction data, and hard-negative strategies vary (Table 1). **(B)** Derivation of the SFM architecture from the Boltzmann distribution governing molecular binding at thermal equilibrium. The probability that agent residue *i* contacts target residue *j* follows the Boltzmann distribution (top). Representing residue properties as feature vectors: query vectors **q**_i_ for the agent and key vectors **k**_*j*_ for the target and substituting the dot-product compatibility score 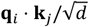 into the Boltzmann equation recovers the softmax attention mechanism of Vaswani et al^34^. The value vector **v**_*j*_ encodes the information content of target residue *j* (chemical identity, charge, shape) that is transferred upon contact; the output **o**_i_ is the contact-probability-weighted sum of value vectors across all target residues. The final matrix equation **O** = softmax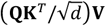 is the standard transformer attention equation, derived here from the biophysics of molecular binding.

**Table 1.** Dataset sources for SFMs. All datasets are publicly available. Bilinearity: *given* = binding energy admits analytical decomposition from sequence; *emergent* = must be learned from data. The architecture, training objective, and evaluation protocol are identical across all six SFMs.

| SFM | Domain | Data source | Pairs | Unique agents / targets | Encoders (agent / target) | Bilinearity |
| --- | --- | --- | --- | --- | --- | --- |
| **tSFM** | TF–DNA | JASPAR 2022 | 20,540 | 1,060 TFs / 15,425 DNA motifs | ESM-2 / DNABERT-2 | Given |
| **eSFM** | Enzyme–substrate | ReactZyme | 177,442 | 177,389 enzymes / 2,646 substrates | ESM-2 / MoLFormer-XL | Emergent |
| **mhcSFM** | Peptide–MHC I | NetMHCpan 4.1 BA | 168,710 | 38,837 peptides / 105 alleles | ESM-2 / ESM-2 | Given |
| **crisprSFM** | gRNA–genomic OT | CCLMoff | 30,342 | 1,668 guides / 30,308 OT sites | DNABERT-2 / DNABERT-2 | Given |
| **mir-SFM** | miRNA–mRNA | ENCORI CLIP-seq | 206,692 | 135 miRNAs / 110,221 targets | DNABERT-2 / DNABERT-2 | Given |
| **dtSFM** | Drug–target | DAVIS / KIBA / BindingDB | 121,548 | 10,338 drugs / 1,358 targets | MoLFormer-XL / ESM-2 | Emergent |

Each SFM was initiated with a single natural-language specification (**Supplementary Note S1**) provided to an AI coding agent (Claude Code; Anthropic), with the published CALM codebase^3^ as the reference implementation. The coding agent handled all data downloads, file-format parsing, database API calls, sequence embedding, preprocessing pipeline construction, training-script generation, SLURM job submission, and evaluation — tasks that constitute the most labor-intensive and technically demanding component of any computational biology project and that required no manual coding at any stage. The domain expert’s contributions were the scientific decisions enumerated in each specification: the choice of ReactZyme over BRENDA for enzyme–substrate data, the use of HLA binding-groove pseudo-sequences rather than full MHC protein sequences for mhcSFM, the design of within-guide true-versus-false off-target hard negatives for crisprSFM. These are decisions that require domain expertise, not programming skill. Each SFM has approximately 5–7 million trainable parameters and trains in under one hour on a single GPU. Every numerical claim reported in this paper was independently verified by an orthogonal AI auditor (Codex; OpenAI) operating on committed repository artifacts (see Methods; **Supplementary Note S2**).

### SFMs achieve bidirectional cross-modal retrieval across molecular recognition domains

All six SFMs were trained exclusively on publicly available paired-interaction data (**Table 1**): JASPAR^35^ for transcription factor–DNA binding, ReactZyme^36^ for enzyme–substrate catalysis, the NetMHCpan 4.1 binding-affinity training corpus for peptide–MHC^13^, CCLMoff for CRISPR off-targets^37^, ENCORI CLIP-seq for miRNA–mRNA targets^38^, and DAVIS/KIBA/BindingDB^39–41^ for drug–target binding. Pretrained sequence encoders — ESM-2 ^42^, DNABERT-2 ^43^, MoLFormer-X^44^ — were obtained from public model repositories. The architecture was applied unchanged across all six domains; identical optimizer, scheduler, batch size, and evaluation protocol were used throughout (**Supplementary Table S2**).

All six SFMs produce meaningful bidirectional retrieval from the identical architecture (**Table 2**). In the agent-to-target direction, in-distribution R@1 ranges from 27.7% (dtSFM, drug→target) to 98.0% (mir-SFM, miRNA→target). In the target-to-agent direction, in-distribution R@1 ranges from 25.4% (mir-SFM, target→miRNA) to 93.3% (mhcSFM, allele→peptide) (**Fig. 2A**). The asymmetry between directions is biologically expected: mhcSFM’s allele→peptide retrieval (93.4%) exceeds peptide→allele (64.8%) because each HLA allele groove constrains its peptide repertoire to a well-defined binding motif, while individual peptides are short and present degenerate binding features across alleles. dtSFM’s target→drug direction (60.5%) exceeds drug→target (27.7%) because protein targets with annotated ligands occupy a structured region of embedding space. Full R@1, R@5, and R@10 values in both directions are in **Table 2**; per-fold results across all clustering thresholds are in **Supplementary Table S1**.

**Table 2.**
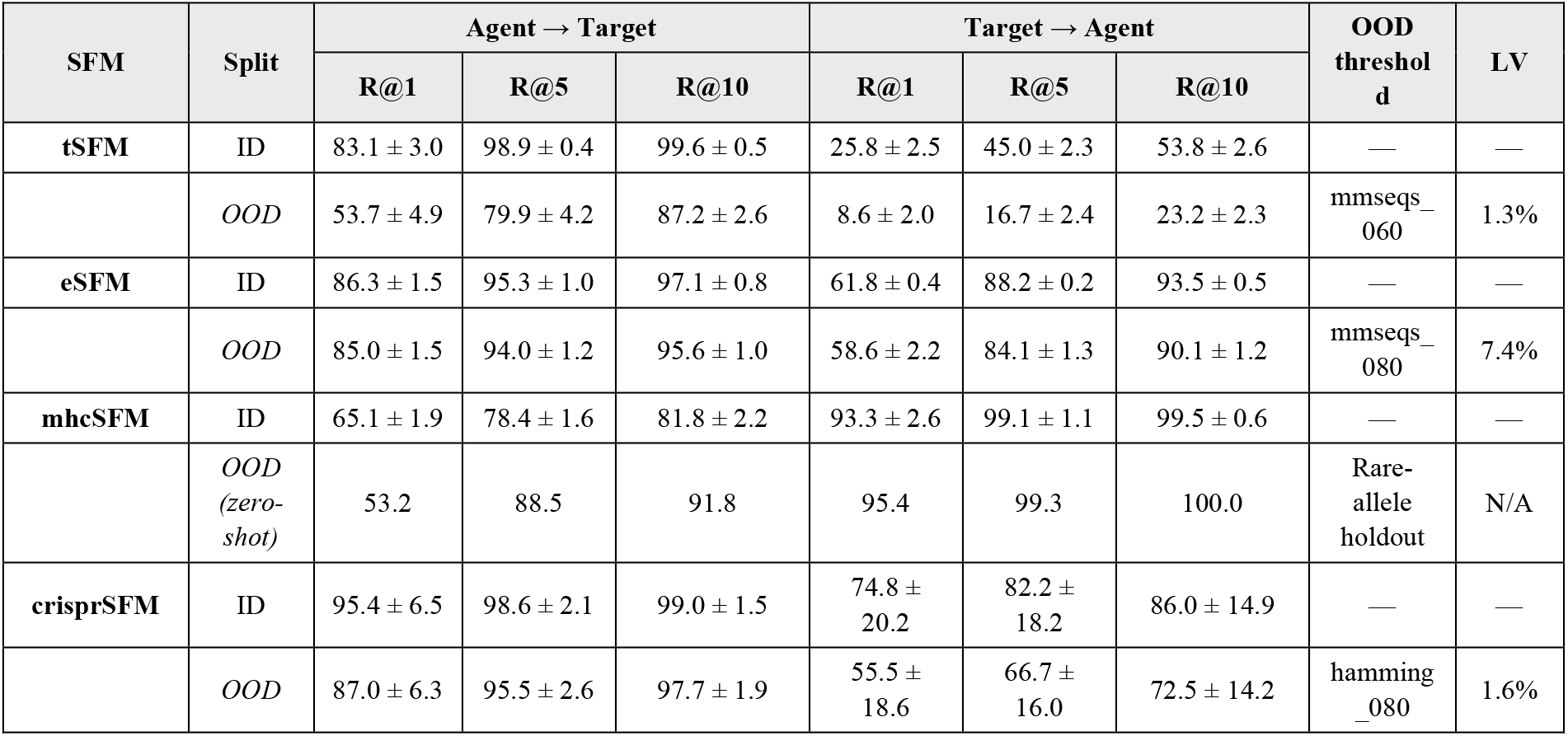
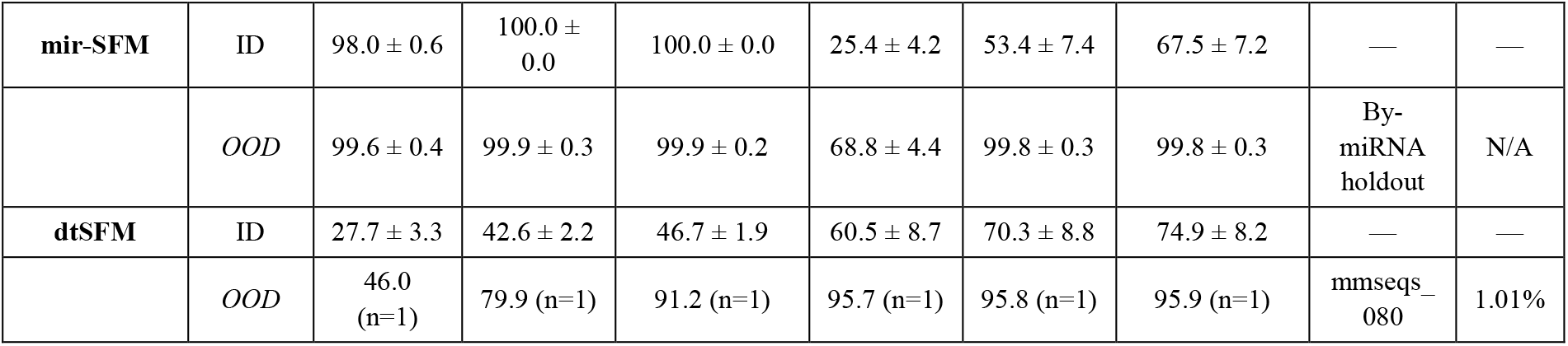
Bidirectional pool-512 retrieval performance across SFMs. Values are cross-fold mean ± s.d. (folds 0–3 excluding fold 4 where the validation set was degenerate; 5-fold where all folds trained successfully). *ID* = in-distribution (identity_100). *OOD* = strictest split-quality threshold passing leakage verification (LV < 10%). Single-fold values are marked (n=1). dtSFM OOD: fold 0 only (folds 1–4 lack checkpoints). mhcSFM OOD: 14 rare alleles held out entirely from training (MMseqs splits fail LV at all thresholds). mir-SFM OOD: by-miRNA holdout. All values are audit-verified.

| SFM | Split | Agent → Target |  |  | Target → Agent |  |  | OOD threshold | LV |
| --- | --- | --- | --- | --- | --- | --- | --- | --- | --- |
|  |  | R@1 | R@5 | R@10 | R@1 | R@5 | R@10 |  |  |
| tSFM | ID | 83.1 ± 3.0 | 98.9 ± 0.4 | 99.6 ± 0.5 | 25.8 ± 2.5 | 45.0 ± 2.3 | 53.8 ± 2.6 | — | — |
|  | OOD | 53.7 ± 4.9 | 79.9 ± 4.2 | 87.2 ± 2.6 | 8.6 ± 2.0 | 16.7 ± 2.4 | 23.2 ± 2.3 | mmseqs_060 | 1.3% |
| eSFM | ID | 86.3 ± 1.5 | 95.3 ± 1.0 | 97.1 ± 0.8 | 61.8 ± 0.4 | 88.2 ± 0.2 | 93.5 ± 0.5 | — | — |
|  | OOD | 85.0 ± 1.5 | 94.0 ± 1.2 | 95.6 ± 1.0 | 58.6 ± 2.2 | 84.1 ± 1.3 | 90.1 ± 1.2 | mmseqs_080 | 7.4% |
| mhcSFM | ID | 65.1 ± 1.9 | 78.4 ± 1.6 | 81.8 ± 2.2 | 93.3 ± 2.6 | 99.1 ± 1.1 | 99.5 ± 0.6 | — | — |
|  | OOD (zero-shot) | 53.2 | 88.5 | 91.8 | 95.4 | 99.3 | 100.0 | Rare-allele holdout | N/A |
| crisprSFM | ID | 95.4 ± 6.5 | 98.6 ± 2.1 | 99.0 ± 1.5 | 74.8 ± 20.2 | 82.2 ± 18.2 | 86.0 ± 14.9 | — | — |
|  | OOD | 87.0 ± 6.3 | 95.5 ± 2.6 | 97.7 ± 1.9 | 55.5 ± 18.6 | 66.7 ± 16.0 | 72.5 ± 14.2 | hamming_080 | 1.6% |
| **mir-SFM** | ID | 98.0 ± 0.6 | 100.0 ± 0.0 | 100.0 ± 0.0 | 25.4 ± 4.2 | 53.4 ± 7.4 | 67.5 ± 7.2 | — | — |
|  | OOD | 99.6 ± 0.4 | 99.9 ± 0.3 | 99.9 ± 0.2 | 68.8 ± 4.4 | 99.8 ± 0.3 | 99.8 ± 0.3 | By-miRNA holdout | N/A |
| **dtSFM** | ID | 27.7 ± 3.3 | 42.6 ± 2.2 | 46.7 ± 1.9 | 60.5 ± 8.7 | 70.3 ± 8.8 | 74.9 ± 8.2 | — | — |
|  | OOD | 46.0 (n=1) | 79.9 (n=1) | 91.2 (n=1) | 95.7 (n=1) | 95.8 (n=1) | 95.9 (n=1) | mmseqs-080 | 1.01% |

**Figure 2.**
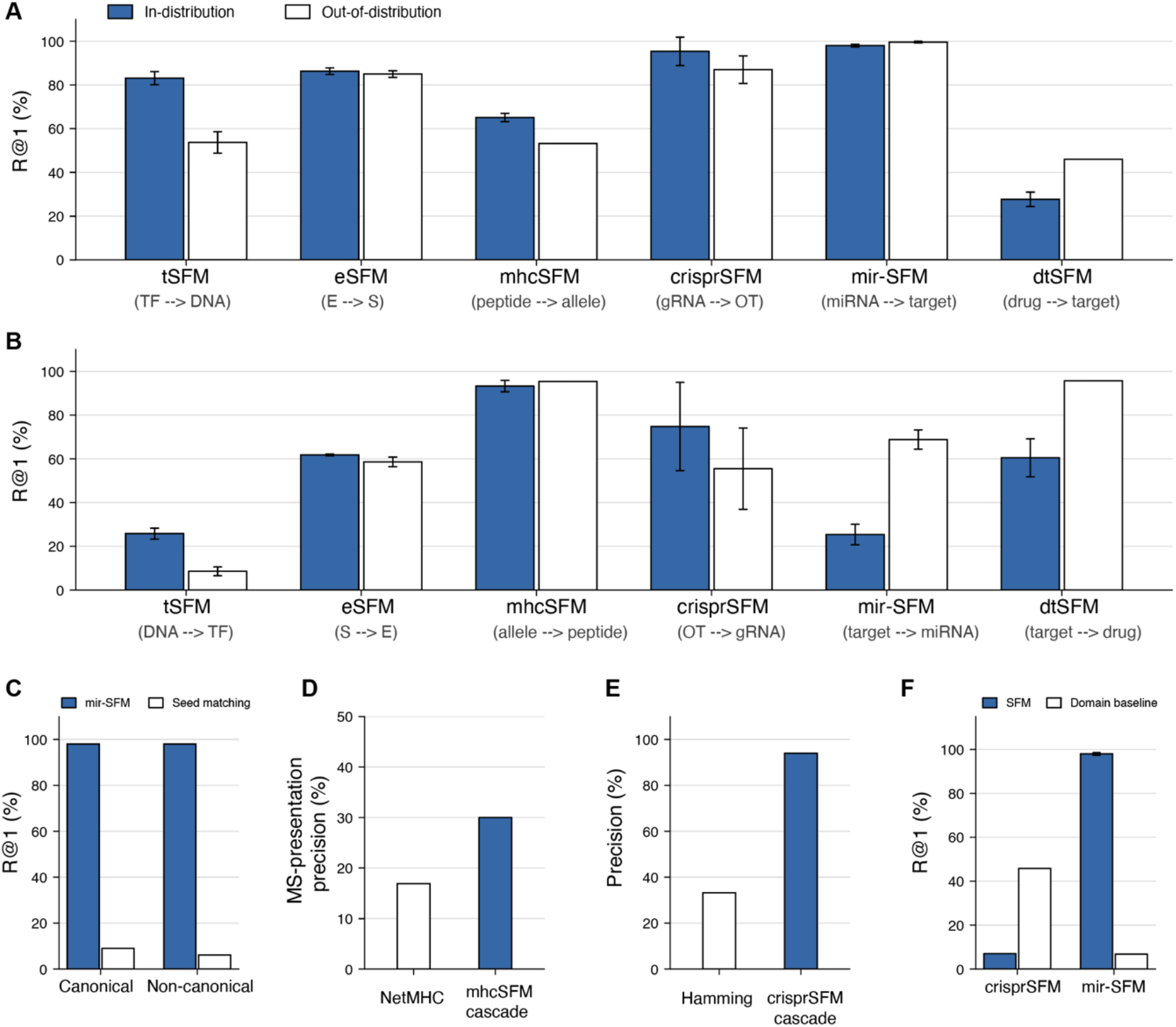
Bidirectional retrieval and external validation across SFMs. All six SFMs share an identical dual-encoder, symmetric InfoNCE architecture; only the encoders, paired training data, and hard-negative strategy differ. **(A)** Agent-to-target R@1 from pools of 512 candidates (R@1 = 0.2% random; 100 random resamplings per fold). Blue bars, in-distribution (identity_100); white bars, best out-of-distribution threshold passing leakage verification (<10%). OOD thresholds: tSFM mmseqs_060, eSFM mmseqs_080, crisprSFM hamming_080, dtSFM mmseqs_080; mhcSFM uses a 14-allele zero-shot holdout; mir-SFM uses a by-miRNA holdout. **(B)** Target-to-agent R@1 (inverse direction), same evaluation. Error bars ± s.d. across folds; bars without error bars denote single-fold evaluations. Fold 4 excluded from means where validation-set degeneracy was detected by the orthogonal audit. **(C)** mir-SFM versus seed-matching baseline, stratified by canonical and non- canonical miRNA–target interactions. **(D)** MS-presentation precision on the Gurung et al.^33^ cancer neoantigen benchmark (86 MS-validated peptide–HLA pairs): NetMHCpan 4.1 EL alone versus NetMHCpan ∩ mhcSFM cascade. **(E)** CRISPR off-target precision at ≤4 mismatches on the CRISPRoffT held-out benchmark (802 candidate sites): Hamming distance alone versus Hamming → crisprSFM re-ranking cascade. **(F)** SFM R@1 versus domain-specific rule-based baseline R@1 for crisprSFM (Hamming distance) and mir-SFM (seed-matching).

OOD performance was evaluated at each similarity-clustering threshold and validated by direct leakage verification (LV; **Table 3**; see below). Four of six SFMs retain meaningful OOD retrieval at the strictest LV-validated threshold: eSFM R@1 = 85.0% at 80% protein identity; tSFM R@1 = 53.7% at 60% identity; crisprSFM R@1 = 87.0% at 80% Hamming identity; dtSFM R@1 = 46.0% at 80% protein identity (single fold). mhcSFM’s similarity-based OOD splits fail LV at all thresholds — 105 alleles encoded as 34-residue pseudo-sequences do not admit strict sequence-identity partitioning — and its OOD generalization is demonstrated by zero-shot retrieval on 14 rare HLA alleles held out entirely from training (R@1 = 95.4%, allele→peptide). mir-SFM’s 135 unique miRNAs produce singleton clusters at all MMseqs2 thresholds; its identity_100 split is a by-miRNA holdout and is inherently OOD with respect to miRNA identity.

**Table 3.**
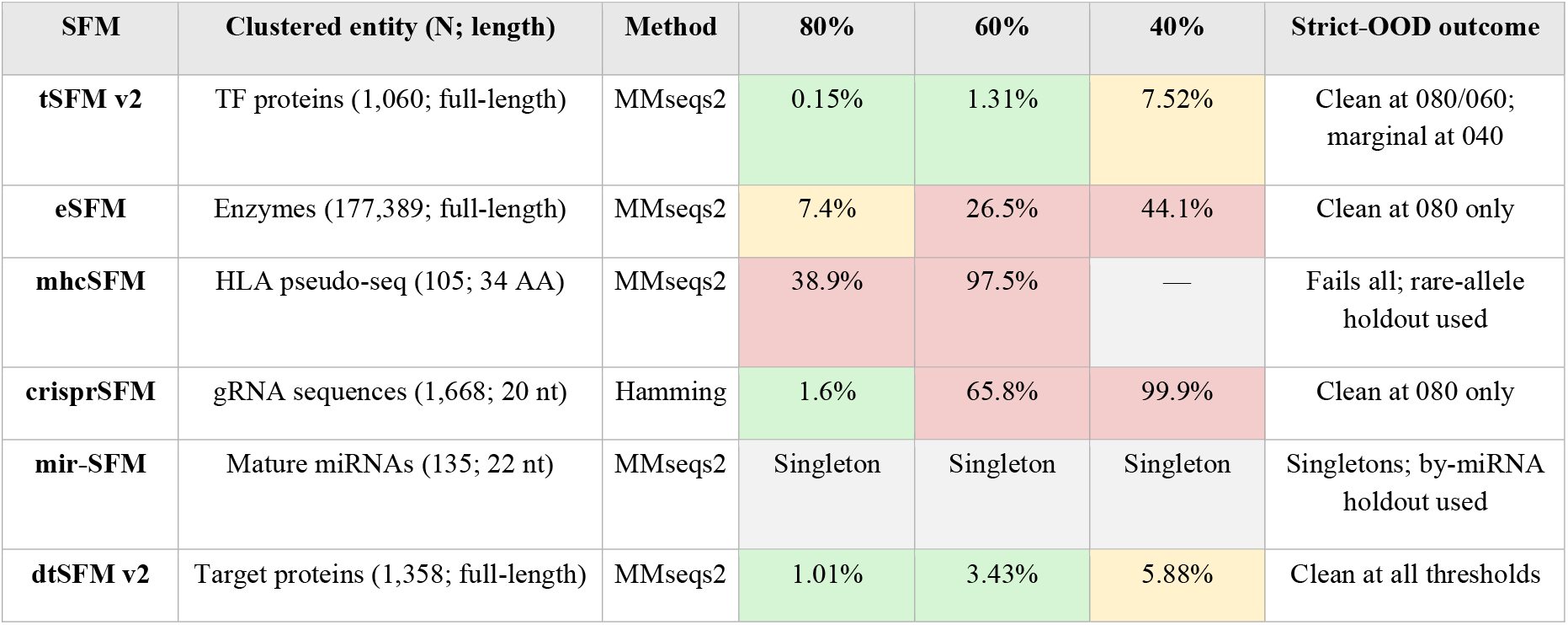
Leakage verification across six SFMs. Leakage = fraction of test entities whose nearest training neighbor exceeds the nominal identity threshold. Cell shading: **green** ≤ 5%; **yellow** 5–10%; **red** > 10% (dropped from claims). Domains with full-length proteins and high entity counts admit strict-OOD splits; small-N or short-input domains require alternative validation.

The dtSFM exhibits a pattern predicted by the MRC framework but absent from standard supervised learning: OOD retrieval exceeds in-distribution retrieval in both directions (drug→target: R@1 = 46.0% OOD vs 27.7% ID; target→drug: R@1 95.7% OOD vs 60.5% ID). This is consistent with the MRC framework’s prediction that when the SFM architecture implements the governing equation^2^, OOD splits remove homology-exploitable shortcuts and regularize the model toward the physical solution.

### SFMs augment standard computational tools

Three external validation experiments demonstrate that SFMs provide operational value when deployed alongside established computational and experimental pipelines (**Fig. 2C-F**).

#### miRNA–mRNA target retrieval

mir-SFM retrieves miRNA targets at 98.0 ± 0.6% R@1 against a seed-matching baseline of 6.8% (**Fig. 2C**). Stratified by target type, seed-matching achieves 9.0% R@1 on canonical-seed targets (21% of the dataset) and 6.1% on non-canonical targets (79%); mir- SFM retrieves both classes at ≥97.5% R@1. Non-canonical miRNA–target interactions constitute approximately 80% of experimentally validated binding events identified by AGO-CLIP-seq^10,32^. Seed-matching — the operating principle of TargetScan^14^ — is architecturally incapable of identifying non-canonical targets because no seed complementarity exists to detect. mir-SFM is the first computational tool that retrieves both canonical and non-canonical miRNA–target interactions from a shared embedding space.

#### Peptide–MHC cascade filter

NetMHCpan 4.1^13^ is the gold-standard predictor for peptide–MHC class I binding and is used in most personalized cancer vaccine pipelines. However, a well-documented limitation is the high false-positive rate of predicted epitopes: binary binding predictions carry approximately 63% false-positive rate against TR-FRET biochemical binding and approximately 85% against mass-spectrometry presentation^33^, a problem recognized by the NetMHCpan developers as arising from training on binding affinity data that models only the single event of peptide–MHC binding rather than the full antigen presentation pathway^45^. In Gurung et al.^33^, 86 cancer neoantigens across 15 prevalent HLA class I alleles were validated by mass-spectrometry immunopeptidomics, providing ground-truth MS-presentation labels. We used this dataset to test whether mhcSFM retrieval-based re-ranking, applied after NetMHCpan 4.1 EL prediction, reduces the false-positive load. The cascade filter — intersecting the NetMHCpan EL top-50 candidates with the mhcSFM retrieval top-50 — nearly doubles MS-presentation precision from 16.9% (NetMHCpan alone) to 30.0% (cascade) (**Fig. 2D**), halving the false-positive rate on candidate vaccine peptide selection.

#### CRISPR off-target cascade filter

Computational CRISPR off-target prediction begins with enumerating all genomic sites within a specified mismatch threshold of the gRNA. At the clinically standard threshold of ≤4 mismatches, a single gRNA typically returns hundreds to thousands of candidate off-target sites, the vast majority of which are false positives — sites that match the sequence but are not cleaved in vivo^7^. The current pipeline uses Hamming distance (or Cas-OFFinder)^15^ as the initial pre-filter, followed by experimental validation via GUIDE-seq or CIRCLE-seq and related methods^46^. On the CRISPRoffT held-out benchmark^47^ (115 guides absent from training, 802 candidate off-target sites at ≤4 mismatches, 266 true and 536 false), Hamming distance alone achieves 33.2% precision at the top-50 cutoff. A cascade filter pairing Hamming pre-filtering with crisprSFM re-ranking raises precision to 94.0% (**Fig. 2E**) — a 2.8-fold reduction in the experimental validation burden.

#### Mirror-image test of the MRC framework

The crisprSFM and mir-SFM results constitute a mirror-image test of the MRC framework’s central empirical prediction (**Fig. 2F**). In CRISPR off-target recognition, binding is dominated by Watson-Crick complementarity and Hamming distance is near-optimal — the SFM does not improve upon it as a standalone (R@1 = 10.2% vs R@1 = 45.8%). In miRNA target recognition, binding involves seed complementarity, non-canonical pairing, and context-dependent rules that sequence matching cannot capture — the SFM exceeds seed-matching by 91 percentage points. The MRC framework predicts that contrastive embeddings add value where binding rules are emergent and complex, and that rule-based baselines only suffice where binding follows canonical complementarity. Both predictions are confirmed by the data.

### Physics-derived SFM architecture exhibits training dynamics consistent with parameter estimation

Across the six domains, SFM training exhibits dynamics that differ from standard supervised deep learning in ways predicted by the MRC framework for models performing parameter estimation in a known governing equation rather than function approximation in an unknown function class (**Fig. 3**).

**Figure 3.**
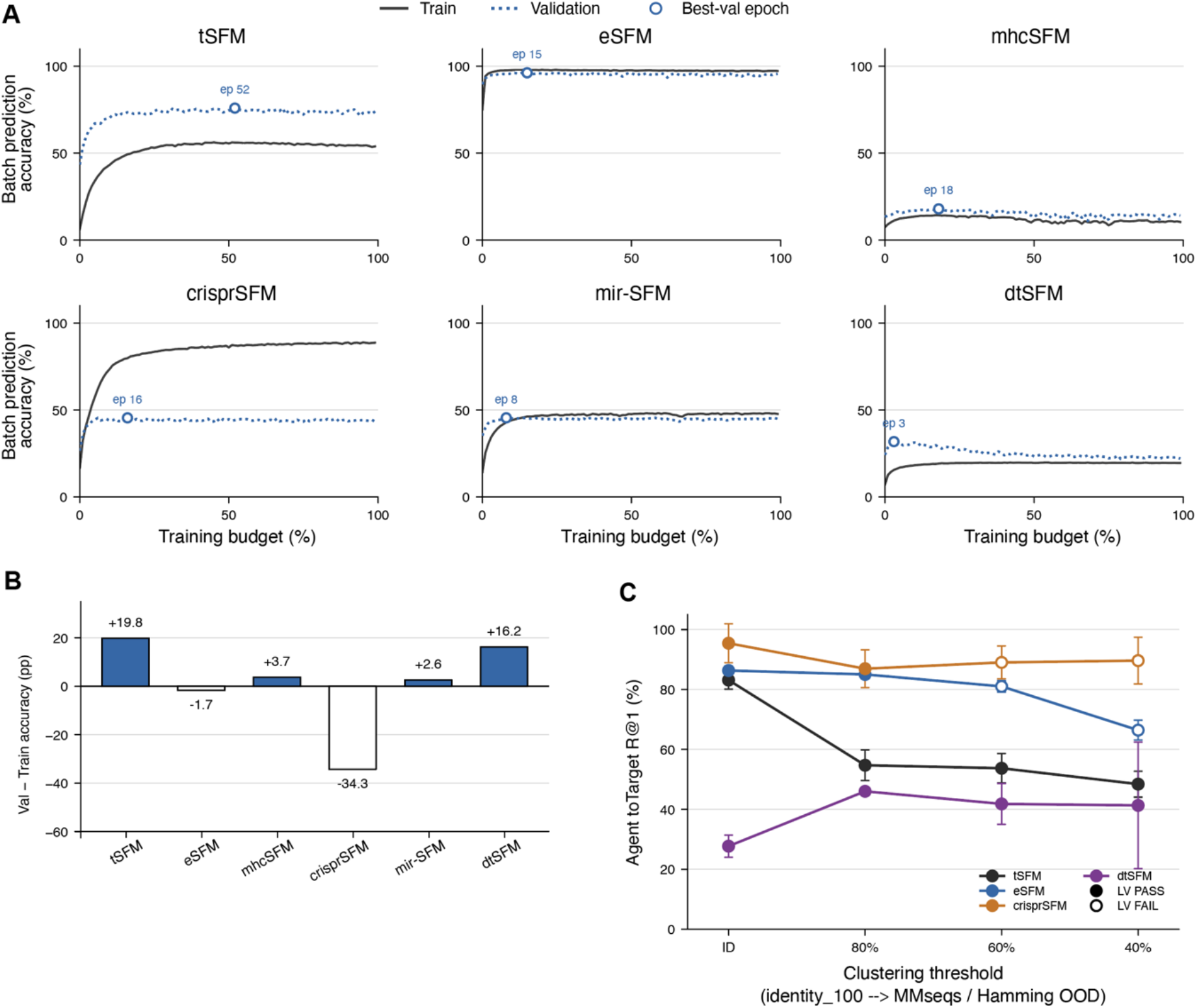
Training dynamics across six SFMs. (A) Training (solid grey) and validation (dotted blue) batch prediction accuracy versus training budget (% of 100 epochs) for each SFM, fold 0, identity_100 split. Batch prediction accuracy is the symmetric InfoNCE batch-64 accuracy averaged over both contrastive directions; absolute heights reflect both model performance and the number of unique partners per batch. All six models were trained with identical settings (AdamW, lr = 10^-3^, CosineAnnealingWarmRestarts T0 = 20, batch size 64). Open blue circles mark the best-validation epoch. **(B)** Validation- minus-training accuracy (percentage points) at the best-validation epoch. Positive bars indicate validation exceeded training at checkpoint selection; negative bars indicate the conventional pattern. **(C)** Agent-to-target pool-512 R@1 as a function of out-of-distribution clustering threshold for four SFMs with multiple evaluated thresholds. Thresholds use MMseqs2 protein sequence identity (tSFM, eSFM, dtSFM) or Hamming nucleotide identity (crisprSFM). Filled markers, leakage verification passed (LV < 10%); open markers, LV failed (shown for reference, excluded from paper claims). mhcSFM and mir-SFM are omitted because similarity-based splits are degenerate in both domains. Error bars ± s.d. across folds where available; points without error bars are single-fold evaluations

In four of six domains (eSFM, mhcSFM, mir-SFM, dtSFM), validation accuracy peaks within the first 3–18% of the training budget (**Fig. 3A**). dtSFM converges fastest, reaching its best checkpoint at epoch 3 of 100; mir-SFM at epoch 8; eSFM at epoch 15; mhcSFM at epoch 18. The two remaining domains (tSFM, crisprSFM) converge later. Both share a common feature: a competing trivial sequence-similarity baseline exists in their loss landscape (PWM-decomposable binding for tSFM; Watson-Crick complementarity for crisprSFM), which the SFM must match before the contrastive embedding provides additional discriminative value.

At the best-checkpoint epoch, four of six SFMs show an inverted train-test gap: validation accuracy exceeds training accuracy (**Fig. 3B**). The gap is +19.8%for tSFM, +16.2% for dtSFM, +3.7% for mhcSFM, and +2.6% for mir-SFM. crisprSFM shows the conventional pattern (−34.3%); eSFM is near zero (−1.7%). The inverted gap — in which a model generalizes before it memorizes — runs counter to the standard bias-variance description of supervised learning^48^, where training accuracy upper-bounds validation accuracy. Under the MRC framework, this pattern is predicted when the SFM architecture implements the governing equation: the physically correct solution is reached before the model has capacity to overfit training-specific patterns.

As OOD threshold stringency increases, retrieval performance decays monotonically but does not collapse to chance in any domain (**Fig. 3C**). tSFM, eSFM, and crisprSFM show gradual decay across threshold levels. dtSFM exhibits the inverse: OOD retrieval (R@1 = 46.7% at 80% protein identity) exceeds in-distribution retrieval (R@1 = 27.7%), consistent with the MRC framework’s prediction that OOD splits act as regularizers when the SFM architecture implements the governing equation.

These three patterns — fast convergence, inverted train-test gap, and OOD non-degradation — co-occur in four of six domains. Their joint occurrence has not been reported in standard supervised deep learning. They are consistent with the prediction that SFM training is parameter estimation in a known governing equation and provide the empirical basis for systematic scaling experiments testing this prediction across data-diversity gradients.

### Leakage verification for strict out-of-distribution evaluation

The standard approach to out-of-distribution evaluation in computational biology uses sequence-similarity clustering (typically MMseqs2^49^) to partition data into train and test sets at a nominal identity threshold. The assumption is that no test-set entity shares more than the specified identity with any training-set entity. This assumption is incorrect for greedy clustering algorithms: MMseqs2 uses a greedy incremental algorithm that assigns each sequence to the first cluster whose representative exceeds the threshold. Because the threshold is applied only to the cluster representative (not to all members), two sequences in different clusters can share identity far exceeding the nominal threshold. The degree of leakage depends on the structure of the input space — alphabet size, sequence length, and entity count jointly determine how often the greedy heuristic produces inter-cluster identity violations.

We measured leakage directly for all six SFMs at all evaluated thresholds by computing pairwise identity between every test-set entity and every training-set entity (**Table 3**). The results reveal a systematic pattern. Domains with full-length protein sequences and thousands of unique entities (tSFM: 1,060 TFs; eSFM: 177,389 enzymes; dtSFM: 1,358 targets) achieve low leakage (0.15–7.4% at 80% threshold). Domains with short input sequences or small entity counts show moderate to catastrophic leakage: crisprSFM (20-nt gRNAs, 1,668 guides) achieves 1.6% at 80% but 65.8% at 60%; mhcSFM (34-AA pseudo-sequences, 105 alleles) shows 38.9% at 80%; mir-SFM (22-nt miRNAs, 135 entities) produces singleton clusters at all thresholds. OOD feasibility via similarity clustering depends on input diversity, not on the clustering protocol alone.

The orthogonal AI verification framework (see Methods) independently confirmed these leakage measurements across all six SFMs. In several cases, the auditor identified leakage at thresholds that the initial analysis had assumed to be clean. All OOD claims reported in this paper are restricted to thresholds where directly measured leakage falls below 10% (**Table 3**); thresholds exceeding this bound are dropped from paper claims regardless of nominal clustering identity.

## DISCUSSION

The six SFMs reported here demonstrate that the architectural prescription derived from the convergence equation^1,2^ transfers across molecular recognition domains without modification. The same dual-encoder, symmetric InfoNCE architecture — with identical training configuration, evaluation protocol, and loss function — produces meaningful cross-modal retrieval in domains spanning protein–DNA, protein–small-molecule, protein–peptide, nucleotide–nucleotide, and nucleotide–RNA binding. The cross-domain transfer is itself the strongest available test of the MRC framework: if the convergence equation was not the governing equation of molecular binding, or if the five architecture conditions did not generalize, a single architecture applied unchanged across six domains would not produce six working models — it would produce six failures requiring domain-specific architectural rescue. That vibe coding by a domain expert with no programming experience sufficed to build all six SFMs is a direct consequence of the architectural prescription: when the governing equation specifies the architecture, the ML engineering load is removed, and the binding constraint becomes scientific judgment. The interface-density analysis (**Supplementary Table S3**) provides a practical design principle for future SFM construction: mean pooling is sufficient when at least one side of the interaction is interface-dense; domains where both sides are dilute require domain-informed preprocessing (such antibody-antigen binding)^3^ or architectural extensions. The orthogonal AI verification protocol — in which an independent AI agent audits all numerical claims from committed artifacts — provides a replicable standard for vibe-coded scientific software.

Several limitations apply. These are prototype models trained on a fraction of the paired-interaction data available in each domain. No prospective wet-lab validation has been performed; the external validations reported use held-out computational benchmarks, not new experimental data. Standalone SFM performance does not exceed established gold-standard tools in all cases — crisprSFM loses to Hamming distance as a standalone classifier, and mhcSFM is less accurate than NetMHCpan 4.1 as a standalone binding predictor — though in both cases the cascade-filter configuration provides substantial operational value. The training-dynamics patterns (fast convergence, inverted train-test gap, OOD non-degradation) are observed in four of six domains; the two exceptions (crisprSFM, tSFM) have identified structural explanations and that the 4/6 pattern is not universal.

The computational requirements for SFM construction are minimal. Each SFM has approximately 5–7 million trainable parameters — the two feedforward projection heads that map frozen encoder outputs into the shared embedding space. The pretrained encoders (ESM-2, DNABERT-2, MoLFormer-XL) are frozen during training; only the projection heads are learned. With precomputed embeddings, training a single fold completes in approximately one hour on a single consumer-grade GPU. The full cross-validation protocol (four OOD thresholds × five folds = 20 runs) completes overnight on a university cluster. No multi-GPU training, no distributed computing, no specialized hardware is required. Combined with the open-source release of all code, checkpoints, datasets, and evaluation pipelines, this places SFM construction within reach of any research group with access to a single GPU and public paired-interaction data. The cross-attentive decoder extension we have previously specified^2,3^ converts every retrieval SFM into a bidirectional generative model: given a target, generate candidate binding agents; given an agent, predict its target repertoire. Production-grade SFMs across the six domains reported here — trained on the full available data, extended with decoders, and validated prospectively — would constitute a physics-derived, sequence-native computational layer for molecular recognition, operating across the binding modalities that organize biology and medicine.

## Methods

### Model architecture

Each SFM consists of two frozen pretrained encoders, two trainable feedforward projection heads, and a symmetric contrastive loss. The agent encoder *f*_*a*_ maps an agent sequence to a fixed-dimensional embedding; the target encoder *f*_*t*_ does the same for the target sequence. Encoder outputs are mean-pooled across sequence length and projected through a two-layer feedforward network (FFN) with ReLU activation into a shared *d*-dimensional embedding space (*d* = 512 for all six SFMs):

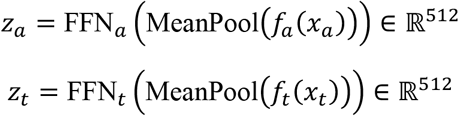

The FFN projection maps from the encoder’s native dimension (*d*_ESM-2_ = 1280, *d*_DNABERT-2_ = 768, *d*_MoLFormer_ = 768) to the shared 512-dimensional space. Encoder weights are frozen during training; only the FFN parameters are learned.

Binding compatibility between agent *x*_a_ and target *x*_t_ is computed as cosine similarity in the shared space:

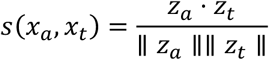

Training minimizes the symmetric InfoNCE loss^50^ with a learned temperature *τ*:

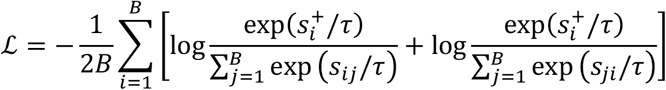

where *B* is the batch size, *s*^+^ is the cosine similarity of the *i*-th positive (matched) pair, and *s*_*ij*_ is the similarity between agent *i* and target *j*. The first term sums over agent-to-target retrieval; the second over target-to-agent. In-batch negatives serve as the denominator: each non-matched pair within the batch acts as a negative example. The temperature *τ* is a learnable scalar initialized to 0.07 with maximum scale 100, corresponding to physical *k*_*B*_*T* in the MRC framework^2^.

This architecture is prescribed by the convergence of softmax attention and the Boltzmann distribution and the five architecture conditions^1,2^ and is identical to the contrastive component of CALM^3^. No architecture search, loss-function ablation, or hyperparameter optimization was performed across domains — the same configuration was applied to all six SFMs.

### Encoder selection

Encoders were selected from publicly available pretrained models to match the molecular modality of each input:

#### Protein sequences

(TF proteins in tSFM, enzymes in eSFM, HLA pseudo-sequences in mhcSFM, drug-target proteins in dtSFM): ESM-2 with 650M parameters, 1280-dimensional output^42^.

#### Nucleotide sequences

(DNA motifs in tSFM, gRNA and genomic off-targets in crisprSFM, miRNA and mRNA target sites in mir-SFM): DNABERT-2, 768-dimensional output^43^. For mir-SFM, RNA sequences were converted to DNA alphabet (U→T) prior to encoding.

#### Small-molecule SMILES

(drug compounds in dtSFM, enzyme substrates in eSFM): MoLFormer-XL, 768-dimensional output^44^.

All encoders were obtained from the Hugging Face model hub. No encoder fine-tuning was performed; encoders remain frozen throughout training.

### Training configuration

All six SFMs were trained with the following shared configuration: AdamW optimizer^51^ with learning rate 1 × 10^−3^ and weight decay 0.2; CosineAnnealingWarmRestarts scheduler with *T*_0_ = 20 and *T*_mult_ = 1; batch size 64; 100 training epochs (tSFM used 200 epochs due to an early configuration that was not subsequently standardized). Model selection used the best validation-set accuracy checkpoint. Unique sequences were embedded once and cached; the training loop operated on precomputed embedding indices, eliminating redundant encoder forward passes for sequences appearing in multiple pairs. Each SFM has approximately 5–7 million trainable parameters (two FFN projection heads with hidden dimension 2048); pretrained encoders remain frozen. Training a single fold completes in minutes to approximately one hour on a single GPU with precomputed embeddings.

### Cross-validation and data splitting

Each SFM was evaluated under a five-fold cross-validation scheme. For each fold, the dataset was split into train (60%), validation (20%), and test (20%) partitions. Four OOD regimes were evaluated per SFM where applicable: identity_100 (in-distribution, random split), and similarity-clustered splits at 80%, 60%, and 40% identity thresholds. Protein-sequence clustering used MMseqs2^49^ with –min- seq-id at the target threshold. Nucleotide-sequence clustering used Hamming distance. Entire clusters were assigned to a single partition to prevent inter-partition homology leakage at the nominal threshold. Fold 4 was excluded from all reported means due to a cross-validation split degeneracy (see Orthogonal AI Verification).

### Retrieval evaluation

Retrieval performance was evaluated using pool-512. For each test-set pair (*x*_a_, *x*_*t*_), the true target *x*_t_ was placed into a pool of 512 candidates (1 positive + 511 randomly sampled negatives from the test set). The agent embedding *z*_a_ was compared to all 512 candidate embeddings by cosine similarity, and candidates were ranked. Recall at rank *k* (R@*k*) measures the fraction of test pairs for which the true target appears in the top-*k* ranked candidates. R@1, R@5, and R@10 are reported. The procedure was repeated for 100 random pool compositions and averaged. Evaluation was performed bidirectionally: agent→target (given an agent, retrieve its target) and target→agent (given a target, retrieve its agent). Random baseline for pool-512 is 1/512 ≈ 0.2%.

### Leakage verification

Nominal similarity-clustering thresholds do not guarantee that all inter-partition sequence identities fall below the threshold. MMseqs2 uses a greedy incremental algorithm that evaluates identity only against the cluster representative, not against all cluster members. To verify OOD claims, we performed direct pairwise leakage measurement: for each test-set entity, the maximum sequence identity to any training-set entity was computed. The leakage fraction *λ* is the proportion of test entities for which this maximum exceeds the nominal threshold. OOD claims are reported only for thresholds where *λ* < 0.10 (**Table 3**). For thresholds where *λ* ≥ 0.10, a conservative filtered retrieval metric is computed: *R*_filtered_ = (*R*_raw_ − *λ*)/(1 − *λ*), bounding worst-case performance if all leaked entities were trivially correct. Full leakage measurements are in **Supplementary Table S1**.

### Vibe coding protocol

All six SFMs were developed using a vibe coding workflow: a domain expert (S.T.R.) with biological and ML conceptual knowledge but no Python programming experience directed an AI coding assistant (Claude Code; Anthropic) via natural language prompts to implement data preprocessing, training, and evaluation pipelines. The CALM codebase^3^ served as the reference implementation; each SFM was instantiated by specifying the data source, encoder pair, hard-negative strategy, and evaluation configuration in a natural-language session prompt. The AI agent generated all code, SLURM job scripts, and analysis pipelines. A representative session-initiating prompt is provided in **Supplementary Note S1**. The domain expert’s contributions were scientific decisions: selection of training data and filtering criteria, encoder choice, hard-negative design (e.g., cofactor-vs-substrate decision for eSFM, within-guide true-vs-false off-targets for crisprSFM), evaluation design, and biological interpretation. No code was written by hand.

### Orthogonal AI verification

Every numerical claim was independently verified by a second AI agent (Codex; OpenAI) operating on artifacts committed to a version-controlled repository prior to audit initiation. The auditing agent had access to the committed codebase, training logs, evaluation outputs, and configurations — but not to the development conversation or the developing agent’s context. For each SFM, the auditor executed a three-leg verification: (i) reproduction of every paper-reportable number from archived source artifacts, classified as PASS, PARTIAL, or FAIL; (ii) direct pairwise leakage measurement across all OOD splits; (iii) computation of conservative filtered retrieval metrics under worst-case leakage assumptions. The audit identified and resolved a cross-validation split degeneracy affecting fold 4 in all six SFMs, a first-author citation misattribution (mhcSFM), an arithmetic discrepancy between writeup and source data (mhcSFM), and a writeup digit imprecision (mir-SFM). Zero unresolved FAIL verdicts remain across all six SFMs. Complete audit records are in **Supplementary Files S1–S6**. The full protocol is described in **Supplementary Note S2**.

## Data and code availability

All training data were derived from publicly available databases (see **Table 1**). Each SFM has 5–7 million trainable parameters and can be trained in under one hour on a single GPU. Pre-trained model weights for all six SFMs are available on Hugging Face: https://huggingface.co/SFM-BIIE-ETHZ. Code, training and evaluation pipelines, and orthogonal audit records are available on Github: https://github.com/Reddy-BIIE-ETHZ/Vibe-Coding-SFMs.

## Acknowledgments

We would like to acknowledge ETH Zurich Scientific IT Services for High Performance Computing and Swiss National Supercomputing Centre (CSCS).

---

**Supplementary Table S1.** Complete pool-512 retrieval results across six SFMs. R@1, R@5, and R@10 (in %) for each SFM across all evaluated clustering thresholds, all cross-validation folds, and both retrieval directions. Values from 100-trial pool-512 evaluation. Fold 4 is shown but excluded from reported means where the cross-validation split was degenerate (validation set empty or epoch-0 checkpoint); asterisks (^*^) mark such folds. Leakage verification leakage rates for each threshold are reported in main-text Table 3. Agent → Target is the forward biological query (given an agent, retrieve its binding target); Target → Agent is the inverse. eSFM per-fold values are not archived; only CL- verified cross-fold means are shown.

| Threshold | Fold | Agent → Target |  |  | Target → Agent |  |  |
| --- | --- | --- | --- | --- | --- | --- | --- |
|  |  | R@1 | R@5 | R@10 | R@1 | R@5 | R@10 |
| identity_100 | 0 | 81.5 | 98.8 | 99.9 | 27.5 | 45.6 | 52.1 |
|  | 1 | 84.6 | 98.7 | 99.5 | 27.4 | 47.1 | 56.5 |
|  | 2 | 79.9 | 98.5 | 99.0 | 22.3 | 41.6 | 51.2 |
|  | 3 | 86.5 | 99.6 | 100.0 | 26.1 | 45.8 | 55.5 |
|  | 4 * | 27.8 | 66.5 | 81.0 | 4.5 | 14.1 | 22.0 |
|  | Mean ± SD | 83.1 ± 3.0 | 98.9 ± 0.4 | 99.6 ± 0.5 | 25.8 ± 2.5 | 45.0 ± 2.3 | 53.8 ± 2.6 |
| mmseqs_080 | 0 | 62.2 | 86.7 | 91.5 | 13.8 | 24.5 | 32.4 |
|  | 1 | 52.7 | 74.6 | 85.2 | 11.9 | 20.1 | 28.1 |
|  | 2 | 48.5 | 78.8 | 87.1 | 7.9 | 12.4 | 20.0 |
|  | 3 | 53.5 | 80.7 | 86.1 | 7.4 | 13.6 | 21.8 |
|  | 4 * | 56.5 | 74.6 | 80.8 | 10.1 | 18.0 | 25.2 |
|  | Mean ± SD | 54.7 ± 5.1 | 79.1 ± 5.0 | 86.1 ± 3.8 | 10.2 ± 2.7 | 17.7 ± 4.9 | 25.5 ± 5.0 |
| mmseqs_060 | 0 | 49.7 | 74.7 | 83.3 | 9.2 | 16.0 | 22.4 |
|  | 1 | 49.0 | 79.9 | 87.2 | 5.5 | 13.9 | 20.7 |
|  | 2 | 52.3 | 77.3 | 86.5 | 8.7 | 16.0 | 22.3 |
|  | 3 | 56.4 | 82.4 | 89.0 | 8.7 | 17.6 | 24.0 |
|  | 4 * | 60.8 | 85.3 | 90.0 | 11.2 | 20.3 | 26.7 |
|  | Mean ± SD | 53.7 ± 4.9 | 79.9 ± 4.2 | 87.2 ± 2.6 | 8.6 ± 2.0 | 16.7 ± 2.4 | 23.2 ± 2.3 |
| mmseqs_040 | 0 | 52.7 | 76.0 | 84.9 | 9.7 | 18.9 | 25.5 |
|  | 1 | 44.2 | 74.8 | 83.7 | 6.6 | 14.6 | 22.2 |
|  | 2 | 50.7 | 77.4 | 88.4 | 5.5 | 10.9 | 17.7 |
|  | 3 | 51.0 | 77.6 | 88.3 | 8.1 | 13.3 | 19.1 |
|  | 4 * | 43.3 | 70.8 | 82.4 | 7.6 | 15.1 | 21.2 |
|  | Mean ± SD | 48.4 ± 4.3 | 75.3 ± 2.8 | 85.5 ± 2.7 | 7.5 ± 1.6 | 14.6 ± 2.9 | 21.1 ± 3.0 |

**eSFM** (*E*→*S* forward / *S*→*E* inverse)
| Threshold | Fold | Agent → Target |  |  | Target → Agent |  |  |
| --- | --- | --- | --- | --- | --- | --- | --- |
|  |  | R@1 | R@5 | R@10 | R@1 | R@5 | R@10 |
| identity_100 | 0 | N/A | N/A | N/A | N/A | N/A | N/A |
|  | 1 | N/A | N/A | N/A | N/A | N/A | N/A |
|  | 2 | N/A | N/A | N/A | N/A | N/A | N/A |
|  | 3 | N/A | N/A | N/A | N/A | N/A | N/A |
|  | 4 | N/A | N/A | N/A | N/A | N/A | N/A |

| | Mean ± SD | 86.3 ± 1.5 | 95.3 ± 1.0 | 97.1 ± 0.8 | 61.8 ± 0.4 | 88.2 ± 0.2 | 93.5 ± 0.5 |
| --- | --- | --- | --- | --- | --- | --- | --- |
| mmseqs_080 | 0 | N/A | N/A | N/A | N/A | N/A | N/A |
|  | 1 | N/A | N/A | N/A | N/A | N/A | N/A |
|  | 2 | N/A | N/A | N/A | N/A | N/A | N/A |
|  | 3 | N/A | N/A | N/A | N/A | N/A | N/A |
|  | 4 | N/A | N/A | N/A | N/A | N/A | N/A |
| | Mean ± SD | 85.0 ± 1.5 | 94.0 ± 1.2 | 95.6 ± 1.0 | 58.6 ± 2.2 | 84.1 ± 1.3 | 90.1 ± 1.2 |
| mmseqs_060 | 0 | N/A | N/A | N/A | N/A | N/A | N/A |
|  | 1 | N/A | N/A | N/A | N/A | N/A | N/A |
|  | 2 | N/A | N/A | N/A | N/A | N/A | N/A |
|  | 3 | N/A | N/A | N/A | N/A | N/A | N/A |
|  | 4 | N/A | N/A | N/A | N/A | N/A | N/A |
| | Mean ± SD | 81.0 ± 1.9 | 92.1 ± 1.6 | 94.6 ± 1.0 | 53.9 ± 2.5 | 79.7 ± 3.0 | 86.7 ± 2.4 |
| mmseqs_040 | 0 | N/A | N/A | N/A | N/A | N/A | N/A |
|  | 1 | N/A | N/A | N/A | N/A | N/A | N/A |
|  | 2 | N/A | N/A | N/A | N/A | N/A | N/A |
|  | 3 | N/A | N/A | N/A | N/A | N/A | N/A |
|  | 4 | N/A | N/A | N/A | N/A | N/A | N/A |
| | Mean ± SD | 66.4 ± 3.3 | 79.2 ± 2.1 | 83.4 ± 1.9 | 40.6 ± 3.0 | 64.0 ± 3.4 | 73.0 ± 3.0 |

**mhcSFM** (peptide→allele forward / allele→peptide inverse)
| Threshold | Fold | Agent → Target |  |  | Target → Agent |  |  |
| --- | --- | --- | --- | --- | --- | --- | --- |
|  |  | R@1 | R@5 | R@10 | R@1 | R@5 | R@10 |
| identity_100 | 0 | 64.6 | 77.0 | 79.8 | 95.0 | 99.4 | 99.5 |
|  | 1 | 65.9 | 78.8 | 81.8 | 94.5 | 99.9 | 100.0 |
|  | 2 | 63.3 | 76.6 | 80.6 | 93.4 | 99.2 | 99.5 |
|  | 3 | 68.0 | 80.2 | 85.4 | 95.0 | 99.7 | 100.0 |
|  | 4 * | 63.9 | 79.6 | 81.3 | 88.8 | 97.2 | 98.6 |
| | Mean ± SD | 65.1 ± 1.9 | 78.4 ± 1.6 | 81.8 ± 2.2 | 93.3 ± 2.6 | 99.1 ± 1.1 | 99.5 ± 0.6 |
| mmseqs_080 | 0 | 71.5 | 94.0 | 94.6 | 97.2 | 98.3 | 99.2 |
|  | 1 | 68.3 | 91.2 | 91.9 | 94.7 | 98.9 | 100.0 |
|  | 2 | 54.9 | 82.6 | 82.6 | 90.4 | 94.9 | 96.7 |
|  | 3 | 67.5 | 96.8 | 97.3 | 99.3 | 100.0 | 100.0 |
|  | 4 * | 77.4 | 97.5 | 98.0 | 99.1 | 100.0 | 100.0 |
| | Mean ± SD | 67.9 ± 8.3 | 92.4 ± 6.0 | 92.9 ± 6.2 | 96.1 ± 3.7 | 98.4 ± 2.1 | 99.2 ± 1.4 |
| mmseqs_060 | 0 | 75.9 | 97.4 | 97.6 | 98.0 | 98.9 | 99.0 |
|  | 1 | 62.5 | 86.6 | 87.2 | 91.3 | 96.6 | 99.3 |
|  | 2 | 74.3 | 99.5 | 99.5 | 96.1 | 97.9 | 98.8 |
|  | 3 | 71.9 | 98.5 | 98.5 | 99.7 | 100.0 | 100.0 |
|  | 4 * | 61.6 | 86.9 | 87.3 | 92.0 | 97.4 | 99.6 |
| | Mean ± SD | 69.2 ± 6.7 | 93.8 ± 6.5 | 94.0 ± 6.2 | 95.4 ± 3.7 | 98.2 ± 1.3 | 99.3 ± 0.5 |
| zero-shot rare alleles | 0 | 53.2 | 88.5 | 91.8 | 95.4 | 99.3 | 100.0 |
|  | 1 | N/A | N/A | N/A | N/A | N/A | N/A |
|  | 2 | N/A | N/A | N/A | N/A | N/A | N/A |
|  | 3 | N/A | N/A | N/A | N/A | N/A | N/A |
|  | 4 | N/A | N/A | N/A | N/A | N/A | N/A |
| | Mean ± SD | 53.2 (n=1) | 88.5 (n=1) | 91.8 (n=1) | 95.4 (n=1) | 99.3 (n=1) | 100.0 (n=1) |

**crisprSFM** (*gRNA*→*OT* forward / *OT*→*gRNA* inverse)
| Threshold | Fold | Agent → Target |  |  | Target → Agent |  |  |
| --- | --- | --- | --- | --- | --- | --- | --- |
|  |  | R@1 | R@5 | R@10 | R@1 | R@5 | R@10 |
| identity_100 | 0 | 98.6 | 99.7 | 100.0 | 87.5 | 92.2 | 94.6 |
|  | 1 | 99.0 | 99.5 | 99.5 | 87.7 | 94.2 | 95.5 |
|  | 2 | 98.8 | 99.8 | 100.0 | 82.8 | 90.5 | 93.1 |
|  | 3 | 98.1 | 99.5 | 99.5 | 81.3 | 88.2 | 90.5 |
|  | 4 * | 82.4 | 94.4 | 96.0 | 34.6 | 46.0 | 56.4 |
| | Mean ± SD | 95.4 ± 6.5 | 98.6 ± 2.1 | 99.0 ± 1.5 | 74.8 ± 20.2 | 82.2 ± 18.2 | 86.0 ± 14.9 |
| hamming_080 | 0 | 93.8 | 98.2 | 99.1 | 73.1 | 84.6 | 89.0 |
|  | 1 | 90.9 | 96.0 | 98.6 | 79.2 | 83.3 | 85.1 |
|  | 2 | 86.8 | 96.6 | 98.6 | 45.9 | 58.3 | 65.3 |
|  | 3 | 87.9 | 96.3 | 98.3 | 51.1 | 65.0 | 73.3 |
|  | 4 * | 75.4 | 90.5 | 94.0 | 28.2 | 42.1 | 49.7 |
| | Mean ± SD | 87.0 ± 6.3 | 95.5 ± 2.6 | 97.7 ± 1.9 | 55.5 ± 18.6 | 66.7 ± 16.0 | 72.5 ± 14.2 |
| hamming_060 | 0 | 91.3 | 98.6 | 99.7 | 63.8 | 79.2 | 85.6 |
|  | 1 | 93.9 | 99.6 | 100.0 | 67.9 | 80.5 | 85.8 |
|  | 2 | 93.2 | 97.5 | 97.8 | 70.4 | 80.8 | 84.5 |
|  | 3 | 87.8 | 96.7 | 98.1 | 53.4 | 67.7 | 73.5 |
|  | 4 * | 79.0 | 92.6 | 95.9 | 32.6 | 44.9 | 54.2 |
| | Mean ± SD | 89.0 ± 5.4 | 97.0 ± 2.4 | 98.3 ± 1.5 | 57.6 ± 13.8 | 70.6 ± 13.7 | 76.7 ± 12.2 |
| hamming_040 | 0 | 94.4 | 98.8 | 99.5 | 74.2 | 87.1 | 89.7 |
|  | 1 | 91.3 | 95.1 | 96.7 | 71.5 | 80.9 | 83.7 |
|  | 2 | 97.2 | 99.5 | 99.5 | 73.2 | 86.2 | 92.2 |
|  | 3 | 90.4 | 97.8 | 99.0 | 71.1 | 80.3 | 84.0 |
|  | 4 * | 74.8 | 88.6 | 92.8 | 31.6 | 40.3 | 48.7 |
| | Mean ± SD | 89.6 ± 7.8 | 96.0 ± 4.0 | 97.5 ± 2.6 | 64.3 ± 16.4 | 75.0 ± 17.5 | 79.7 ± 15.8 |

**mir-SFM** (*miRNA*→*target* forward / *target*→*miRNA* inverse)
| Threshold | Fold | Agent → Target |  |  | Target → Agent |  |  |
| --- | --- | --- | --- | --- | --- | --- | --- |
|  |  | R@1 | R@5 | R@10 | R@1 | R@5 | R@10 |
| identity_100 | 0 | 98.4 | 100.0 | 100.0 | 26.4 | 56.3 | 70.9 |
|  | 1 | 98.1 | 100.0 | 100.0 | 26.4 | 56.0 | 70.5 |
|  | 2 | 98.5 | 100.0 | 100.0 | 27.2 | 57.6 | 71.5 |
|  | 3 | 98.3 | 100.0 | 100.0 | 29.5 | 58.3 | 71.5 |
|  | 4 * | 96.8 | 100.0 | 100.0 | 17.3 | 38.6 | 53.0 |
|  | **Mean ± SD** | **98.0 ± 0.6** | **100.0 ± 0.0** | **100.0 ± 0.0** | **25.4 ± 4.2** | **53.4 ± 7.4** | **67.5 ± 7.2** |
| **mmseqs_080** | 0 | 98.8 | 99.4 | 99.5 | 64.0 | 99.3 | 99.3 |
|  | 1 | 99.8 | 100.0 | 100.0 | 67.2 | 99.8 | 99.8 |
|  | 2 | 99.9 | 100.0 | 100.0 | 76.4 | 100.0 | 100.0 |
|  | 3 | 99.7 | 100.0 | 100.0 | 65.7 | 100.0 | 100.0 |
|  | 4 * | 99.8 | 100.0 | 100.0 | 70.7 | 100.0 | 100.0 |
|  | **Mean ± SD** | **99.6 ± 0.4** | **99.9 ± 0.3** | **99.9 ± 0.2** | **68.8 ± 4.4** | **99.8 ± 0.3** | **99.8 ± 0.3** |

**dtSFM** (*drug→target forward / target→drug inverse*)
| Threshold | Fold | Agent → Target |  |  | Target → Agent |  |  |
| --- | --- | --- | --- | --- | --- | --- | --- |
|  |  | R@1 | R@5 | R@10 | R@1 | R@5 | R@10 |
| **identity_100** | 0 | 23.3 | 40.5 | 45.4 | 49.5 | 58.5 | 63.5 |
|  | 1 | 25.3 | 42.3 | 47.6 | 50.9 | 61.0 | 66.9 |
|  | 2 | 31.3 | 44.0 | 47.1 | 65.6 | 76.3 | 80.5 |
|  | 3 | 26.9 | 40.1 | 43.9 | 65.8 | 75.7 | 79.7 |
|  | 4 * | 31.6 | 46.0 | 49.5 | 70.8 | 80.1 | 84.2 |
|  | **Mean ± SD** | **27.7 ± 3.3** | **42.6 ± 2.2** | **46.7 ± 1.9** | **60.5 ± 8.7** | **70.3 ± 8.8** | **74.9 ± 8.2** |
| **mmseqs_080** | 0 | 46.0 | 79.9 | 91.2 | 95.7 | 95.8 | 95.9 |
|  | 1 | N/A | N/A | N/A | N/A | N/A | N/A |
|  | 2 | N/A | N/A | N/A | N/A | N/A | N/A |
|  | 3 | N/A | N/A | N/A | N/A | N/A | N/A |
|  | 4 | N/A | N/A | N/A | N/A | N/A | N/A |
|  | **Mean ± SD** | **46.0 (n=1)** | **79.9 (n=1)** | **91.2 (n=1)** | **95.7 (n=1)** | **95.8 (n=1)** | **95.9 (n=1)** |
| **mmseqs_060** | 0 | 49.0 | 66.6 | 92.3 | 96.4 | 97.3 | 97.6 |
|  | 1 | 33.3 | 37.0 | 49.9 | 90.8 | 92.4 | 93.7 |
|  | 2 | 39.9 | 50.3 | 70.5 | 91.1 | 92.1 | 92.9 |
|  | 3 | 44.8 | 54.7 | 64.1 | 91.5 | 92.5 | 92.8 |
|  | 4 * | 18.3 | 27.7 | 33.7 | 2.1 | 9.8 | 16.0 |
|  | **Mean ± SD** | **37.1 ± 10.7** | **47.3 ± 13.6** | **62.1 ± 19.7** | **74.4 ± 36.2** | **76.8 ± 33.6** | **78.6 ± 31.3** |
| **mmseqs_040** | 0 | 31.1 | 36.5 | 45.0 | 76.3 | 79.3 | 79.8 |
|  | 1 | 47.9 | 62.1 | 77.9 | 95.6 | 97.3 | 97.4 |
|  | 2 | 18.9 | 22.7 | 27.2 | 61.5 | 63.4 | 67.4 |
|  | 3 | 67.4 | 96.3 | 100.0 | 97.6 | 97.8 | 98.0 |
|  | 4 * | 8.7 | 15.7 | 20.6 | 2.2 | 6.8 | 10.7 |
|  | **Mean ± SD** | **34.8 ± 20.9** | **46.7 ± 29.5** | **54.1 ± 30.3** | **66.6 ± 34.8** | **68.9 ± 33.6** | **70.7 ± 32.1** |

Notes. \* Fold 4 excluded from reported means due to a cross-validation split degeneracy independently identified across all six SFMs by the orthogonal AI auditor (see Supplementary Note S2). dtSFM mmseqs\_080 is single-fold (folds 1–4 lack checkpoints; see main-text caption). mhcSFM

**Supplementary Table S2.** Training configuration shared across all six SFMs. The SFM architecture is prescribed by the convergence equation and five architecture conditions (Reddy, 2026a, 2026b); the training configuration is identical across domains. The only domain-specific choices are encoder selection (determined by molecular modality), paired-interaction data source, and hard-negative strategy (Table 1). This configuration matches the contrastive training component of CALM^3^.

| Parameter | Value | Notes |
| --- | --- | --- |
| Embedding dimension (d) | 512 | Shared projection space for all SFMs |
| FFN projection | 2-layer, ReLU activation | Maps encoder output to shared space |
| Pooling | Mean pooling | Over sequence length dimension |
| Loss function | Symmetric InfoNCE | Oord et al. (2018); bidirectional |
| Negatives | In-batch | Batch size determines negative count |
| Temperature ($\tau$) | Learned, init = 0.07, max scale = 100 | Corresponds to physical kBT (Reddy, 2026b) |
| Optimizer | AdamW | Loshchilov & Hutter (2019) |
| Learning rate | $1 \times 10^{-3}$ | |
| Weight decay | 0.2 |  |
| Scheduler | CosineAnnealingWarmRestarts | $T_0 = 20$, $T_{mult} = 1$ |
| Batch size | 64 |  |
| Training epochs | 100 | tSFM: 200 (early config, not standardized) |
| Encoder weights | Frozen | Only FFN projection heads are trained |
| Model selection | Best validation accuracy | Checkpoint saved per epoch |
| Cross-validation | 5-fold (60/20/20 train/val/test) | Fold 4 excluded from means |
| Evaluation | Pool-512, 100 random trials | Bidirectional R@1/R@5/R@10 |
| Agent encoders | ESM-2 (1280-d) / DNABERT-2 (768-d) / MoLFormer-XL (768-d) | Selected by molecular modality (Table 1) |
| Target encoders | ESM-2 (1280-d) / DNABERT-2 (768-d) / MoLFormer-XL (768-d) | Selected by molecular modality (Table 1) |
| OOD clustering | MMseqs2 (protein) / Hamming (nucleotide) | Thresholds: 80%, 60%, 40% identity |
| Leakage verification | Direct pairwise measurement | OOD claims require $\lambda < 10\%$ (Table 3) |

**Supplementary Table S3.** Interface density across SFMs. Interface density is the fraction of input sequence positions that participate in the binding interface. Mean pooling aggregates the full encoder output across all positions; when binding-relevant positions are a small fraction of the input, the binding signal is diluted by non-interface sequence. The six SFMs in this paper each have at least one interface-dense side, and mean pooling is sufficient without masking. CALM is the published comparator^3^: both antibody (∼10% paratope on a 230-residue VH-VL) and antigen (∼3–8% epitope) are interface-dilute, requiring epitope/paratope masking at preprocessing. mhcSFM uses an analogous strategy: the 34-residue binding-groove pseudo-sequence is extracted at preprocessing, converting a moderately dilute HLA sequence into a fully dense input. Both are domain-informed preprocessing decisions that reflect biological knowledge of the binding interface, not architectural modifications to the SFM.

| SFM | Agent (length) | Target (length) | Agent interface density | Target interface density | Domain-specific preprocessing | Mean-pool sufficient |
| --- | --- | --- | --- | --- | --- | --- |
| **crisprSFM** | gRNA (20 nt) | Genomic OT (23 nt) | ~100% | ~100% | None | Yes |
| **mir-SFM** | miRNA (22 nt) | mRNA target (30 nt) | ~100% | ~80% | None | Yes |
| **mhcSFM** | Peptide (8–11 AA) | HLA pseudo-seq (34 AA) | ~100% | ~100% | Binding-groove pseudo-sequence extraction | Yes |
| **tSFM** | TF protein (DBD) | DNA motif | ~50% | ~80% | None | Yes |
| **dtSFM** | Drug (SMILES) | Target protein (full) | ~100% | ~5–10% | None | Yes |
| **eSFM** | Substrate (SMILES) | Enzyme (full) | ~100% | ~5% | None | Yes |
| **CALM** | Ab VH+VL (~230 AA) | Antigen (200–1000 AA) | ~10% | ~3–8% | Epitope/paratope masking (5 Å interface) | Required masking |

## Supplementary Note S1: Vibe Coding Protocol for SFM Construction

### Overview

Each Specificity Foundation Model in this paper was initiated with a single natural-language specification provided to an AI coding assistant (Claude Code; Anthropic). The specification contains the domain expert’s scientific decisions — binding modality, data source, encoder choice, hard-negative strategy, validation design — and delegates all implementation to the AI agent. No code was written by hand. The template below shows the modular structure common to all six SFM specifications. The mhcSFM example that follows is the exact prompt that initiated the peptide–MHC SFM development session and produced a working prototype within days.

### Generic template

#### SESSION 1, PROMPT 1

I am [NAME], a [ROLE] at [INSTITUTION]. I am an experimental biologist — I have NO coding experience. Explain every step in plain language and teach me what things mean as we go.

Read Context.md and check memory files.

#### ## What [SFM_NAME] is

[SFM_NAME] predicts [BINDING_MODALITY] specificity. The agent is [AGENT_DESCRIPTIO N].

The target is [TARGET_DESCRIPTION]. Bilinearity is [GIVEN/EMERGENT] because [JUSTIFICATION]. All five architecture conditions are satisfied under [PHYSICAL ASSUMPTION] (Reddy, Methods for Molecular Recognition Computing, bioRxiv, 2026).

#### ## Data source

[DATABASE_NAME] ([CITATION]) contains [N] pairs covering [AGENT_COUNT] unique agents and [TARGET_COUNT] unique targets. Download from [URL].

[DATA FILTERING DECISIONS — what to include, what to exclude, and why.]

#### ## Encoder choice

- Agent encoder: [ENCODER_NAME] ([DIMENSION]-dim) — [JUSTIFICATION]
- Target encoder: [ENCODER_NAME] ([DIMENSION]-dim) — [JUSTIFICATION]

#### ## Training conditions (match all other SFMs)

- Architecture: FFN projection head, mean pooling, d_model=512
- Temperature: learned, initialized tau=0.07 (max_scale=100)
- Training: 100 epochs, batch size 64, AdamW (lr=0.001, wd=0.2)
- Scheduler: CosineAnnealingWarmRestarts (T0=20, Tmult=1)
- Evaluation: pool size 512, 100 random trials, bidirectional R@1/5/10
- OOD splits: [CLUSTERING_METHOD] at 40/60/80% on [WHICH_ENTITY] sequences ## Key domain considerations

[DOMAIN-SPECIFIC DECISIONS — hard negatives, filtering, one-to-many relationships, known pitfalls, biological priors that inform preprocessing.]

#### ## Validation experiment — plan BEFORE training

[VALIDATION STRATEGY — temporal split, holdout entity, external benchmark, head-to-head comparison with gold-standard tool. Justify the choice.]

#### ## Learnings from prior SFMs

[ACCUMULATED KNOWLEDGE — infrastructure notes, known bugs, efficiency tricks, patterns observed in earlier SFMs that inform this one.]

#### ## The 7-step workflow

1. Data assessment
2. Preprocessing and embedding
3. Similarity clustering for OOD splits
4. Smoketest
5. Full training
6. Pool-512 evaluation
7. Validation experiment Let’s start with Step 1.

### Worked example: mhcSFM (peptide–MHC class I specificity)

The following is the exact prompt provided to Claude Code at the start of the mhcSFM development session. It is reproduced verbatim. The resulting model achieved 93.4% R@1 allele→peptide and 64.8% R@1 peptide→allele in-distribution, 95.4% R@1 on 14 held-out rare HLA alleles with zero training data, and a cascade filter with NetMHCpan 4.1 that nearly doubled MS-presentation precision on the Gurung et al. (2023) cancer neoantigen benchmark.

#### SESSION 1, PROMPT 1

I am Sai Reddy, a professor at ETH Zurich. I am an experimental biologist and immunologist — I have NO coding experience. Explain every step in plain language, tell me exactly WHERE to type each command (Mac Terminal vs Euler login node vs Euler GPU job), and teach me what things mean as we go. Every prompt and output in this chat will be published in the “Vibe-Coding SFMs” companion paper.

Read Context.md and check memory files (especially infra_modules.md, workflow_strategy.md).

### What mhcSFM is

mhcSFM predicts peptide–MHC binding specificity. The agent is a short peptide (8-11 amino acids), the target is an MHC class I allele (the groove region, α1–α2 domains). BOTH sides are protein sequences, so both use ESM-2 — architecturally the simplest SFM (same encoder type for agent and target, like crisprSFM used DNABERT-2 for both). Bilinearity is EMERGENT. All five architecture conditions are satisfied (Reddy, 2026b).

This is MY core domain — I am an immunologist. I have deep expertise in peptide-MHC biology, HLA alleles, and immunopeptidomics. I will be making strong domain-expert decisions in this chat.

### This papers feeds into

“Vibe-Coding SFMs” (VC-5Apr.md) — needs full prompt trail + validation experiment

### Why this domain is exciting for the SFM framework

Existing methods (NetMHCpan 4.1, MHCflurry 2.0) are trained on 850K+ data points covering ∼170 alleles — but they do NOT learn a shared metric space. They cannot do zero-shot retrieval for unseen alleles. mhcSFM would be the first contrastive model that maps peptides and MHC alleles to a shared embedding space, enabling: (a) given a peptide, retrieve which alleles present it, (b) given an allele, retrieve which peptides it presents, (c) zero-shot prediction for rare/novel HLA alleles by embedding similarity.

There are >13,000 known HLA class I alleles. Only ∼150 have substantial binding data. mhcSFM’s value is in generalizing to the other 12,850.

#### Data sources (this is where mhcSFM shines — massive data)

PRIMARY: IEDB binding assays — quantitative peptide-MHC binding affinity data (IC50 values). NetMHCpan 4.1 training data includes ∼850K data points. We can download from IEDB directly.

SECONDARY: Mass spectrometry immunopeptidomes — peptides directly identified as presented by specific HLA alleles. Key datasets: - Sarkizova et al. (Nature Biotechnology, 2020): 95 mono- allelic cell lines, ∼186K HLA-I peptides across 92 alleles - Abelin et al. (Immunity, 2017): mono- allelic profiling - IEDB MS-derived ligand data

For the SFM, we need BINARY binding pairs (peptide X binds allele Y). From IEDB: - Binding affinity: IC50 < 500 nM = binder (standard threshold) - Eluted ligand: directly observed by mass spec = binder

Estimated dataset size: 200K-500K peptide-allele pairs across 100-170 alleles. This is the LARGEST dataset of any SFM domain.

#### Training conditions (match tSFM/crisprSFM/dtSFM)

- Architecture: FFN projection head, mean pooling, d_model=512
- Agent encoder: ESM-2 (1280-dim) for peptide — NOTE: peptides are very short (8-11 aa). ESM-2 was designed for full proteins. Check that short peptide embeddings are meaningful.
- Target encoder: ESM-2 (1280-dim) for MHC allele groove region (α1-α2 domains, ∼180 aa)
- Temperature: learned, initialized tau=0.07 (max_scale=100)
- Training: 100 epochs, batch size 64, AdamW (lr=0.001, wd=0.2)
- Scheduler: CosineAnnealingWarmRestarts (T0=20, Tmult=1)
- Checkpoint: best validation accuracy
- Evaluation: pool size 512, 100 random trials, bidirectional R@1/5/10
- OOD splits: MMseqs2 at 40/60/80% on MHC allele sequences + in-distribution

NOTE ON TEMPERATURE: Same learned temperature for cross-domain comparability. Physical temperature is theoretically appropriate (Boltzmann selection at 310K) but deferred.

### Key domain-expert decisions

1. MHC INPUT REPRESENTATION: Use the HLA pseudo-sequence (34 contact residues from the binding groove) rather than the full MHC sequence. This is what NetMHCpan uses — it captures allele-specific groove geometry in a compact representation. Alternatively, use the full α1-α2 domain (∼180 aa). I want to discuss this.
2. OOD CLUSTERING: Cluster by HLA supertype (groups of alleles with similar binding properties, e.g., A02 supertype, B07 supertype). This is the biologically meaningful OOD variable — we want to test whether mhcSFM generalizes to unseen HLA supertypes. MMseqs2 on MHC sequences should approximate supertype groupings.
3. PEPTIDE LENGTH: MHC class I primarily presents 9-mers, but 8-mers and 10-11-mers also bind. Include all lengths. The model should learn length-dependent binding preferences from the data.
4. HARD NEGATIVES: From crisprSFM, we learned that hard negatives dramatically improve discrimination. For mhcSFM, natural hard negatives exist: peptides that bind one allele but NOT a closely related allele (e.g., binds HLA-A*02:01 but not HLA-A*02:07). IEDB has extensive negative data.

### Validation experiment — plan BEFORE training

Candidate approaches: 1. TEMPORAL SPLIT: Train on IEDB data before 2024, validate on data deposited 2024-2026. IEDB tracks deposition dates. 2. RARE ALLELE HOLDOUT: Train on common alleles (A*02:01, B*07:02, etc.), hold out rare alleles with recent MS data. Test zero-shot retrieval. 3. NEW IMMUNOPEPTIDOME: ESCAPE-seq (Borrman et al., Nature Genetics, 2025) screened 75,000+ peptide-HLA combinations. Hold out alleles from this study.

I’m leaning toward option 2 (rare allele holdout) — this directly tests mhcSFM’s unique value proposition: generalizing to alleles with little or no training data.

### Learnings from tSFM, crisprSFM, and dtSFM

- Euler: module load stack/2024-06 gcc/12.2.0 python/3.11.6 + venv (NOT conda)
- Always PYTHONUNBUFFERED=1 in SLURM scripts
- Use /cluster/scratch/for data and output (home quota = 50 GB)
- Embed unique sequences only if dataset has duplicates (index-based loader from data_tsfm.py) — CRITICAL here since ∼170 unique alleles but 500K pairs
- fold_list=“[0]” not outer_fold=0, wandb=false for compute nodes
- Hard negatives via label masking (Option B from crisprSFM) — include non-binding peptide-allele pairs as in-batch negatives
- SLURM 20-job array for full training, ALPS for scaling

### The 7-step workflow

1. Data assessment — download IEDB data, assess size/quality, decide on MHC representation, identify validation holdout
2. Preprocessing — embed peptides and MHC alleles with ESM-2, create metadata + splits
3. MMseqs2 clustering — generate OOD splits at 40/60/80% on MHC allele sequences
4. Smoketest — small test run to verify pipeline
5. Full training — 4 OOD thresholds × 5 folds = 20 runs on Euler
6. Pool-512 evaluation — fill Tables 24-25
7. Scaling experiments — 25 runs on ALPS, fill Tables 32-33 rows

Let’s start with Step 1: Data Assessment. Help me download IEDB peptide-MHC class I binding data and assess what we have. Tell me what to do on my Mac Terminal.

## Supplementary Note S2: Orthogonal AI Verification Protocol

### Purpose

Vibe coding — natural-language-directed software construction using AI coding agents — removes the coding barrier for domain experts building foundation models, but introduces a verification challenge: the developer cannot inspect the generated code with the same fluency as a trained programmer. Orthogonal AI verification addresses this by using a second, independent AI agent to audit the code and claims produced by the first. The protocol described here was developed iteratively across six Specificity Foundation Models and is reported as a replicable standard for future vibe- coded scientific software.

### Protocol

The protocol has seven steps:

#### Step 1 — Develop with Agent A

The domain expert works with an AI coding assistant (in this work, Claude Code; Anthropic) to design, preprocess, train, evaluate, and document each SFM. All development occurs in a version-controlled repository. The expert makes scientific decisions (data sources, encoder selection, hard-negative strategy, evaluation design); the AI agent writes and executes the code.

#### Step 2 — Preserve artifacts before audit

Before initiating the audit, all paper-reportable artifacts are committed to the repository: training logs, evaluation outputs, configuration files, split-index files, and durable summary CSVs. The commit hash marks the frozen state that the auditor will verify against. No artifact created after this commit is admissible as audit evidence.

#### Step 3 — Draft an audit specification

The developer writes a specification enumerating every claim to be verified, organized into two categories: sandbox-auditable (SA) items covering methodology (architecture, hyperparameters, data counts, preprocessing correctness) and claim-level (CL) items covering every numerical value that will appear in the paper. Each item specifies the source artifact, the verification method, and the pass criteria.

#### Step 4 — Launch Agent B in a clean session

A second AI agent (in this work, Codex; OpenAI), operating in a separate session with no access to Agent A’s conversation history, reads the committed repository and the audit specification. Agent B has access to the codebase, data files, logs, and configurations — but not to the development rationale, the natural-language prompts that produced the code, or any context from the development conversation.

#### Step 5 — Agent B executes the three-leg audit

##### Leg 1 (SA + CL verification)

Agent B reads each audit item, locates the source artifact, and either parses the claimed value from logs, recomputes it from committed CSVs, or re-runs analysis scripts. Each item receives a verdict: PASS (value matches within stated tolerance), PARTIAL (value is close but a discrepancy or caveat is documented), or FAIL (value cannot be reproduced or is incorrect).

#### Leg 2 (Leakage verification)

Agent B re-runs the similarity-clustering tool (MMseqs2 for protein-sequence splits, Hamming distance for nucleotide splits) on the committed test and training sets at each nominal out-of-distribution threshold. For each test entity, the auditor measures whether any training entity exceeds the threshold identity. The fraction of test entities that “leak” determines which OOD claims survive into the paper.

#### Leg 3 (Filtered R@1)

For splits where leakage is nonzero, Agent B computes a conservative lower-bound retrieval metric: R_filtered = (R_raw − λ) / (1 − λ), where λ is the measured leakage fraction. This bounds the worst-case performance if all leaked test entities were trivially correct.

#### Step 6 — Review and resolve

The developer reviews Agent B’s deliverables. FAIL verdicts are investigated; some may be re-graded to PARTIAL with documented justification (e.g., when the auditor’s framing does not match the paper’s stated methodology). All re-gradings are recorded in a separate closure-notes document, preserving Agent B’s original verdicts unmodified.

#### Step 7 — Close and tag

The audit deliverables are committed to the repository and a git tag marks closure. The tag message includes the aggregate SA and CL tallies and the paper-level consequences (which OOD claims survive, which are dropped). This tag is the citable reference for the paper’s methodology section.

### What the audit caught

The orthogonal verification framework identified the following issues across six SFMs, all of which were resolved before the claims reported in this paper:

#### Cross-validation split degeneracy (5 of 6 SFMs)

The CALM-inherited nested cross-validation split generator produces a degenerate validation set when the inner fold equals the outer fold, yielding 2–5 validation entries and an untrained best-validation checkpoint. The auditor independently discovered this in eSFM, tSFM, crisprSFM, mhcSFM, and dtSFM. Resolution: fold 4 excluded from all reported means; the detection is automated in the evaluation script.

#### Leakage at nominal OOD thresholds (4 of 6 SFMs)

MMseqs2 greedy clustering at nominal threshold T does not strictly bound pairwise inter-cluster identity. Direct measurement revealed 7.4– 99.9% leakage at lower thresholds across eSFM, crisprSFM, mhcSFM, and dtSFM. Resolution: OOD claims are reported only for thresholds where measured leakage is below 10%; all other thresholds are dropped from paper claims.

#### Citation misattribution (mhcSFM)

The Gurung et al. (2023) *Nature Biotechnology* cancer- neoantigen benchmark was consistently labeled as “Pyke et al. 2024” in development scripts, CSVs, and the writeup — a first-author misattribution. The auditor flagged the discrepancy. Resolution: corrected throughout.

#### Arithmetic discrepancy (mhcSFM)

The writeup stated 5.5% binder removal; recomputation from the source CSV yielded 6.9%. Resolution: corrected to the computed value.

#### Writeup digit imprecision (mir-SFM)

The writeup cited 98.4% R@1 (fold-0 value); the cross-fold mean is 98.0 ± 0.6%. Resolution: corrected to cross-fold mean.

#### Encoder-identifier alias (eSFM)

Preprocessing logs printed “tSFM” labels while processing eSFM data — an inherited debug-label alias. The auditor confirmed data was correct despite the misleading label. Resolution: documented; no impact on results.

### Replicability

The protocol is agent-agnostic. Any combination of AI coding agents can serve as Agent A and Agent B, provided they operate in separate sessions with no shared context. The audit specification format, the three-leg structure, and the PASS/PARTIAL/FAIL classification scheme are described in sufficient detail for any research group to replicate. The complete audit records for all six SFMs are available in the public repository.

